# Disentangling hand and tool processing: distal effects of neuromodulation

**DOI:** 10.1101/2021.12.06.471144

**Authors:** L. Amaral, R. Donato, D. Valério, E. Caparelli-Dáquer, J. Almeida, F. Bergström

## Abstract

The neural processing within a brain region that responds to more than one object category can be separated by looking at the horizontal modulations established by that region, which suggests that local representations can be affected by connections to distal areas, in a category-specific way. Here we first wanted to test whether by applying transcranial direct current stimulation (tDCS) to a region thatre sponds both to hands and tools (posterior middle temporal gyrus; pMTG), while participants performed either a hand- or tool-related training task, we would be able to specifically target the trained category, and thereby dissociate the overlapping neural processing. Second, we wanted to see if these effects were limited to the target area or extended to distal but functionally connected brain areas. After each combined tDCS and training session, participants therefore viewed images of tools, hands, and animals, in an fMRI scanner. Using multivoxel pattern analysis, we found that tDCS stimulation to pMTG indeed improved the classification accuracy between tools vs. animals, but only when combined with a tool training task (not a hand training task). However, surprisingly, tDCS stimulation to pMTG also improved the classification accuracy between hands vs. animals when combined with a tool training task (not a hand training task). Our findings suggest that overlapping but functionally-specific networks can be separated by using a category-specific training task together with tDCS - a strategy that can be applied more broadly to other cognitive domains using tDCS - and demonstrates the importance of horizontal modulations in objectcategory representations.

## Introduction

Object recognition is a complex process that engages different sets of cortical regions. In fact, the way conceptual information is organized in the human brain is still under debate (e.g., Grill-Spector & Malach, 2004; Op de Beeck et al., 2019). Recently, we (and others) have shown that neural processing and the organization of information in one area is dependent not only on local aspects, but also on processes happening within distal but functionally connected regions (Amaral et al., 2021; Lee et al., 2019; Walbrin & Almeida, 2021; see also Almeida et al., 2013; Garcea et al., 2016; Kristensen et al., 2016; Ruttorf et al., 2019). These modulations between areas that belong to a particular domain-specific network – horizontal modulations within a domain – allow for the exchange and integration of different kinds of conceptual information. According to this hypothesis, object topography – that is, the organization of object-related information in the brain – is not only dependent on local computations, but also on connections from distal regions that share a propensity for processing a specific category of objects (Chen et al., 2017; Garcea et al., 2019; see also, Mahon & Caramazza, 2011; Sporns, 2014). Here we will explore these horizontal modulations and focus on the processing of two related categories – hands and tools – as a way of further understanding the organization of object knowledge in the brain.

Early neuroimaging studies suggest that object recognition depends on local neural processes (mainly) within the ventral temporal cortex (VTC) (Grill-Spector & Weiner, 2014; Peelen & Downing, 2017). In fact, several studies have shown that different regions inside VTC present higher BOLD signal change for specific object categories like faces (fusiform face area – FFA, Kanwisher et al., 1997), places (parahippocampal place area - PPA, Epstein & Kanwisher, 1998), bodies (fusiform body area – FBA, Peelen et al., 2005; Schwarzlose et al., 2005), hands (Bracci et al., 2012, 2016) or tools (medial fusiform gyrus - mFUG, Almeida et al., 2013; Chao & Martin, 2000; Garcea & Mahon, 2014) when compared to other high-level categories.

However, recent studies show that these local representations within VTC are (at least partly) shaped by information shared via structural and functional connectivity from distal regions (Hutchison et al., 2014; Saygin et al., 2011, 2016). According to this hypothesis, category-specific representations rely not only on local computations, but also on information processed within distal regions outside VTC that is transferred via long-distance horizontal modulations. As an illustration of this claim, Lee, Mahon and Almeida (Lee et al., 2019) used transcranial Direct Current Stimulation (tDCS) over tool-preferring left inferior parietal areas (IPL), and showed that BOLD signal patterns were modulated by tDCS polarity and, most importantly, that representations in tool-preferring regions within the VTC (specifically, the left medial fusiform gyrus - mFUG), but not in other regions of VTC, changed in a category-specific way (i.e., tDCS changed multivoxel patterns that were elicited by tool stimuli, but not those elicited by face or place stimuli; Lee et al., 2019; see also, Amaral et al., 2021; Chen et al., 2017; Garcea et al., 2019; Walbrin & Almeida, 2021). That is, it is possible to modulate the representational patterns within a specific target region by stimulating a distal area that is functionally connected to, and that shares categorical preferences with that target region. We, therefore, argue that local representations of a specific category can be modulated by information from distal regions that are functionally connected.

One important test to the relevance of distal connectivity and horizontal modulations in conceptual representation and the organization of information in the brain is the situation where a particular region figures critically in the processing of more than one higher-level category (i.e., shows preferential responses to two categories) – can within-domain horizontal modulations disentangle the functionally distinct category-specific networks? Recently, we have demonstrated this to be the case. Specifically, in a fMRI study, we found that functional connectivity from two regions that show an overlap in their preference for tools and hands (left IPL and left pMTG) with other distal areas (e.g., tool or hand preferring regions of the VTC) is differently correlated with categorical preferences: in tool-preferring VTC areas, functional connectivity from left IPL and left pMTG (i.e., the tool/ hand-preferring overlap areas) correlates with local response preferences for tools but not hands, whereas in hand-preferring VTC areas, it correlates with local response preferences for hands but not tools (Amaral et al., 2021). That is, horizontal modulations connecting regions of a domain-specific network (for hands or tools) allow for the separation of the two different networks, despite the overlap response that occurs for both categories in left IPL and left pMTG.

If an overlap response can be separated by focusing on the different domain-specific horizontal modulations, then questions arise as to whether we can enhance this separation effect by biasing the processing towards one of those categories? That is, if we have two categories that both drive responses in a particular overlap region, will enhancing the processing of one of those categories lead to an increase of the category-specific responses elsewhere via horizontal modulations? Here, we address this question by combining tDCS with (cognitive) training tasks to enhance processing for a particular category.

We focused on two functionally related categories – hands and tools (Almeida et al., 2018; Amaral et al., 2021; Bergström et al., 2021; Bracci et al., 2010, 2012). First, we wanted to investigate if tDCS applied to one of the areas where preferences for tool and hand stimuli overlap (i.e., pMTG) would provoke distal effects in other regions of the brain (see Lee et al., 2019; Ruttorf et al., 2019). Second, we wanted to test if we could disentangle the functionally-specific networks for hands and for tools, as these have some overlapping nodes. We did this by applying tDCS to pMTG (or to a control area - medial Prefrontal Cortex; mPFC) in combination with a category-specific training task, prior to an fMRI session.

tDCS is a neuromodulation procedure that adapts neuronal excitability through the depolarization or hyperpolarization of resting membrane potential (Nitsche et al., 2008; Nitsche & Paulus, 2011). Unlike other brain stimulation techniques (e.g., Transcranial Magnetic Stimulation), tDCS does not produce action potentials in the neuronal cell membranes. For this reason, several authors believe that tDCS action relies on the activity already present in the tDCS areas (i.e., before and/or during stimulation) (Stagg & Nitsche, 2011). If tDCS is activity-dependent, we could potentially enhance its effects by triggering a specific cognitive processing prior to (and during) stimulation. That is, we may enhance tDCS effects by cognitively manipulating the network-specific engagement of the system in preparation for tDCS stimulation. As such, in tandem with the tDCS stimulation, we asked the participants to perform an online (pre-MRI) task that focused on one of the categories (hands or tools). By hypothesis, the task will produce task-related neural spiking and tDCS stimulation will be added on top of augmented neural responses.

After the simultaneous high-definition tDCS (to improve focality) and task training session, participants went through an event-related fMRI experiment where we presented images of tools, hands, and animals. The tDCS montages and category tasks were manipulated within participants (such that each participant went through 4 sessions). Using Multivoxel Pattern Analysis (MVPA) over the BOLD patterns for tools or hands from the fMRI session, we showed that stimulating pMTG (combined with the training tasks) leads to different patterns of classification between hands and tools (vs animals).

## Methods

### Participants

Twenty-five subjects participated in this experiment (M = 22 years, SD = 3.5, 8 males). All participants had normal or corrected to normal vision, were right-handed, had no history of neuropsychiatric disorders (e.g., stroke, epilepsy, dementia, depression) or head injury, had no metallic implants, did not intake concurrent medication likely to affect cognition and had no history of alcohol and drug abuse or dependence. Written informed consent was obtained from all participants prior to the beginning of the study. Participants were each paid €40 upon completion of the study. Students from the Faculty of Psychology and Educational Sciences of the University of Coimbra also received course credits for their participation. The study was approved by the Ethical Committee of the Faculty of Psychology and Educational Sciences of the University of Coimbra. Due to signal problems, we excluded data from all runs for one participant. Four participants did not complete all sessions, so we also excluded the data from those participants: two did not finish the experiment, one showed neurological abnormalities, and, for the last subject, the monitor inside the scanner was not working. Consequently, 20 participants (M = 23 years, SD = 3.5, 6 males) were used for the analyses of this study.

### Experimental procedure

All participants included in the analyses completed four sessions with a minimum interval of one week: two sessions using pMTG as the stimulated area and two sessions with mPFC. For each pair of sessions, the training task could be either hand or tool-related (pMTG_hands_, pMTG_tools_, mPFC_hands_, mPFC_tools_). The order of the sessions was randomized and counterbalanced across subjects. Each session always started with the HD-tDCS application and a training task, followed by the fMRI experiment in which participants viewed images of tools, hands, and animals.

### HD-tDCS

We used a battery-driven HD-tDCS system composed of a direct current generator, connected to a HD-tDCS adaptor (Soterix Medical, NY, USA). The electrode montage was planned with the aid of a current flow modeling software which employs finite element method to calculate the resulting electric field in brain regions during the stimulation (HD-Explore - Soterix Medical, NY, USA). The resulting simulation is based on the electrodes’ positions. Electrodes’ deployment was planned in order to optimize focality over the stimulation target area (pMTG), while avoiding the current flow into parietal regions (e.g., inferior parietal lobe - IPL). Since IPL is also known as an overlap area when processing hands and tools, in order to ensure that stimulation targeted the temporal lobe, we only used 3 electrodes (1 anodal centered at TP7, according to the 10-10 EEG system, and two cathodal located 5cm distant from the anodal, see Figure 1). Regarding the mPFC stimulation, we kept the same setup using only 3 electrodes with the anodal located at the Fpz position. The HD-electrodes were placed inside a holder filled with Signa Gel (Parker Laboratories, NJ, USA). Impedance values were examined for each electrode and the intensity of the current was set to 2 mA, delivered for 20 minutes (ramp duration of 1 minute). The tDCS room was immediately adjacent to the MRI scanner, allowing for a fast transfer to the MRI environment right after stimulation. For simplicity, we henceforth refer to it as tDCS.

**Figure 1.**
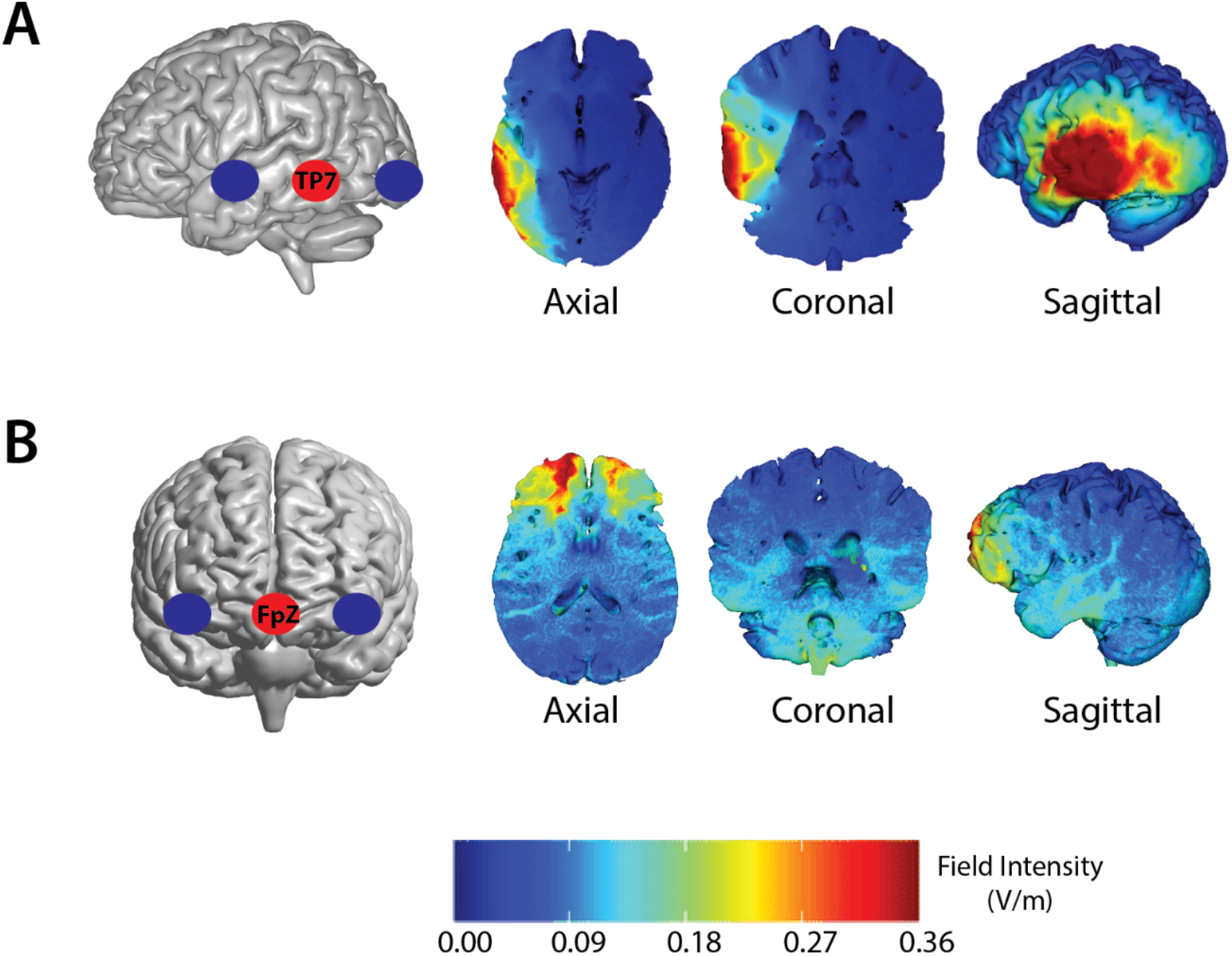
Electrode montage and modeling of the brain current flow. **A)** Electrode montage targeting posterior middle temporal cortex (pMTG) and the respective brain current flow based on HD-explore software (Soterix Medical, NY, USA). **(B)** Electrode montage targeting medial prefrontal cortex (mPFC) and the respective brain current flow showing that this montage did not affect the brain regions included in the pMTG montage.

### Training task

Participants performed two equivalent one-back tasks, one with tool images and another with hand images. All images were black and white, and appeared on the screen for 400ms with a refresh rate of 60 Hz. During the tool training, participants pressed a button when the current and the previous image belonged to a different object (e.g., a glass and a bowl), but not when they belonged to the same (basic level) object, despite potential changes in perspective or exemplars (e.g., different angles or types of a glass). During the hand training, participants pressed a button when the current and previous hand image referred to a different hand side – i.e., right or left hand. Participants saw different perspectives and hand postures.

The responses were collected with a button box (Cedrus Corp.), with their dominant hand (right). We used MATLAB and “A Simple Framework” (ASF; Schwarzbach, 2011) to present stimuli. We measured accuracy and reaction times, and the experiment lasted for 30 minutes (tDCS started at minute 10). Due to a technical problem, the files corresponding to the session pMTG_hands_ in one subject were not saved. Thus, we excluded this participant from our behavioral analysis.

### fMRI task

We used an event-related design with four runs for the fMRI experiment. Participants were presented with centrally fixated gray-scaled images (400*400 pixels) of tools, hands, animals, and places. Each image was presented for 2 sec, followed by a 4 sec fixation period. Participants were asked to detect catch trials (i.e., trials consisting of place images) and press a button every time they saw a place image. The purpose of this task was to keep participants alert while attending to all stimuli. Nonetheless, for all sessions, we used an eye tracker to (subjectively) monitor the individual’s attention (and wakefulness) during the entire task. Stimulus delivery and response collection were controlled using Psychtoolbox (Brainard, 1997) in Matlab (The MathWorks Inc., Natick, MA, USA). Stimuli were presented on an Avotec projector with a refresh rate of 60 Hz and viewed by the participants through a mirror attached to the head coil inside the bore of the MR scanner. Each run began with an 8 sec fixation period and ended with a 16 sec fixation period. Eight different exemplars of tools, hands and animals were used, and each run contained 3 repetitions per stimulus. Six different exemplars of places were used as catch trials and each run contained 1 repetition per each of this stimulus. Due to a technical problem, we were not able to collect the button responses in the first two sessions of subject 1 and 2.

### Data acquisition and statistical analysis

#### Data acquisition

MRI data were acquired using a 3T MAGNETOM Trio whole body MR scanner (Siemens Healthineers, Erlangen, Germany) with a 64-channel head coil. There were four sessions, and each one included four functional runs and one structural scan. Structural MRI data was collected using T1-weighted rapid gradient echo (MPRAGE) sequence (repetition time (TR) = 2530 msec, echo time (TE) = 3.5 msec, slice thickness = 1 mm, flip angle = 7 deg, field of view (FoV) = 256 * 256, matrix size = 256 * 256, bandwidth (BW) = 190 Hz/px, GRAPPA acceleration factor 2). Functional MRI (fMRI) data were acquired using a T2*-weighted gradient echo planar imaging (EPI) sequence (TR = 2000 msec, TE = 30 msec, slice thickness = 3 mm, FoV = 210 * 192, matrix size = 70 * 64, flip angle =75 deg, BW = 2164 Hz/px, GRAPPA acceleration factor 2). Each image volume consisted of 37 contiguous transverse slices recorded in interleaved slice order oriented parallel to the line connecting the anterior commissure to the posterior commissure covering the whole brain.

#### Image preprocessing

We used SPM12 (Wellcome Trust Centre for Neuroimaging, London, UK), run in Matlab R2018b (The MathWorks Inc., Natick, MA, USA), for processing and analysis of structural and functional data. All images were reoriented to approximate MNI space with SPM12 after slice-time correction. The functional data were slice-time corrected to the first slice using a Fourier phase-shift interpolation method, corrected for head motion to the first volume of the first session using 7^th^ degree b-spline interpolation. Structural images were coregistered to the first functional images. Functional data were then normalized to MNI anatomical space using a 12-parameter affine transformation model in DARTEL (Ashburner, 2007) and smoothed with an 8 mm (for ROI localization) and 3 mm (for MVPA) FWHM Gaussian filter.

#### Univariate analysis

For each participant, a fixed-effects analysis was performed by setting up a General Linear Model (GLM) with animals, hands, and tools as regressors of interest; and places (catch trials) as well as motion correction parameters (to covary out signal correlated with head motion) as nuisance regressors. All regressors of interest were convolved with a canonical hemodynamic response function to create the design matrix. Model estimations for each participant were used in a second-level random-effects analysis to account for inter-individual variability.

#### Regions of interest (ROIs)

Two univariate contrasts (tools > animals and hands > animals) were used to select group and individual peak-coordinates for regions engaged by tools and hands. ROIs were defined in two steps, as proposed by Oosterhof and colleagues (Oosterhof et al., 2012). First, we created group-level spheres with 15 mm radius using MarsBaR (Brett et al., 2002) centered on the group’s univariate peak-voxel coordinates. Second, we created individual-level spheres with 15 mm radius centered on each individual’s univariate peak-voxel coordinates but within the group-level spheres.

#### Multivariate pattern analysis

We used a leave-one-run-out cross-validation procedure to train a Support Vector Machine (SVM) classifier to discriminate between z-score normalized beta patterns of two experimental conditions (hands vs. animals OR tools vs. animals). The leave-one-run-out cross-validation procedure ensured that training and testing data was kept completely independent. The multivariate classification analysis was performed with The Decoding Toolbox (Hebart et al., 2015). The group’s average classification accuracies were computed for each condition (i.e., pMTG_hands_, pMTG_tools_, mPFC_hands_, and mPFC_tools_) and for each ROI (i.e., the six hand- and eight tool-related ROIs, separately). ROIs defined by the hands > animals contrast were used to classify hands vs. animals, and ROIs defined by the tools > animals contrast were used to classify tools vs. animals. Thus, we had two different designs depending on the ROIs that were analyzed: (i) 2 (tDCS area: pMTG or mPFC) * 2 (training task: hands or tools) * 6 (hand-ROIs: described in detail in the results section), and (ii) 2 (tDCS area: pMTG or mPFC) * 2 (training task: hands or tools) * 8 (tool-ROIs). The accuracy results were therefore analyzed with a repeated measure ANOVA with these three factors. Specifically, we were interested in whether there was an interaction between the tDCS area and the training task. In addition, we analyzed (for each ROI) the difference in classification accuracy between the pMTG and mPFC conditions. To do this, we compared the classification accuracy between pMTG and mPFC in a paired t-test for each ROI. For the statistical analyses (e.g., ANOVA) we used IBM SPSS Version 22 (IBM Corp., Armonk, NY).

## Results

### Training task

We used a 2 (training task: hands or tools) * 2 (tDCS area: pMTG or mPFC) factorial repeated-measures ANOVA to analyze the accuracy and the reaction times in the training task. Regarding the accuracy values, there was a main effect of the training task (*F*(1,18) = 30.30, *p* < .0001) such that accuracy in the tool task was greater than in the hand task (see Table 1). For the reaction times, we observed the same main effect of the training task (*F*(1,18) = 355.73, *p* < .0001) such that reaction times in the hand task were higher when compared to the tool task (see Table 1).

**Table 1.**
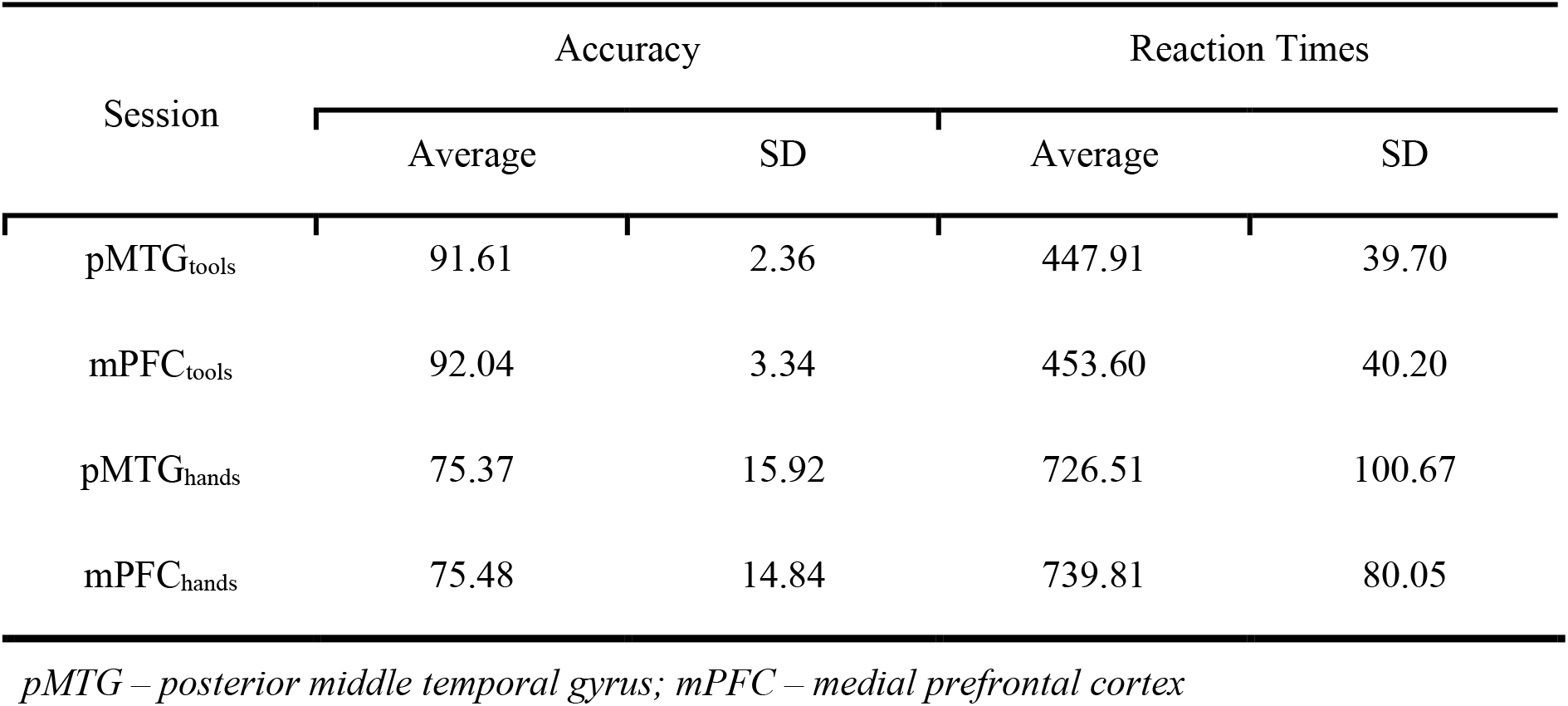
Training task results.

| Session | Accuracy |  | Reaction Times |  |
| --- | --- | --- | --- | --- |
|  | Average | SD | Average | SD |
| pMTG<sub>tools</sub> | 91.61 | 2.36 | 447.91 | 39.70 |
| mPFC<sub>tools</sub> | 92.04 | 3.34 | 453.60 | 40.20 |
| pMTG<sub>hands</sub> | 75.37 | 15.92 | 726.51 | 100.67 |
| mPFC<sub>hands</sub> | 75.48 | 14.84 | 739.81 | 80.05 |

*pMTG – posterior middle temporal gyrus; mPFC – medial prefrontal cortex*

### fMRI results

#### Behavioral task

Participants viewed images of hands, tools, and places inside the scanner, and were instructed to press a button every time they saw an image of a place in order to maintain them awake and attentive to the stimuli. The results show a high hit rate (M = 99%, SD = 1.5) and a low false alarm rate (M = .6%, SD = .7), indicating that participants were, indeed, paying attention to the images.

#### ROI selection

The hands > animals contrast (p<.001, uncorrected) revealed increased BOLD signal change in bilateral posterior parietal cortices (extending across the superior parietal lobe (SPL) and anterior intraparietal sulcus (aIPS), and bilateral pMTG. The tools > animals contrast (p<.001, uncorrected) revealed increased BOLD signal change in the posterior parietal cortices (SPL bilaterally, and extending to aIPS and supramarginal gyrus (SMG) in the left hemisphere), left pMTG, left dorsal occipital cortex (DOC), and medial fusiform gyrus (mFUG). Thus, we chose the following regions as ROIs for hand areas: left SPL (peak t-value = 4.46, MNI coordinates = [−24 −69 60]), right SPL (peak t-value = 5.93, MNI coordinates = [30 −63 60]), left aIPS (peak t-value = 5.23, MNI coordinates = [−30 −39 42]), right aIPS (peak t-value = 4.62, MNI coordinates = [30 −42 48]), left pMTG (peak t-value = 9.37, MNI coordinates = [−51 −66 6]), and right pMTG (peak t-value = 5.42, MNI coordinates = [51 −57 0]). As tool ROIs, we chose left SPL (peak t-value = 6.14, MNI coordinates = [−24 −69 60]), right SPL (peak t-value = 5.87, MNI coordinates = [21 −69 60]), left IPS (peak t-value = 5.62, MNI coordinates = [−24 −57 51]), left SMG (peak t-value = 3.86, MNI coordinates = [−45 −33 39]), left DOC (peak t-value = 6.05, MNI coordinates = [−30 −84 18]), left pMTG (peak t-value = 6.23, MNI coordinates = [−54 −69 −6]), left mFUG (peak t-value = 6.14, MNI coordinates = [−27 −51 −15]), and right mFUG (peak t-value = 4.57, MNI coordinates = [27 −48 −12]).

#### MVPA results

We used two factorial repeated-measure ANOVAs to analyze the accuracy values from our classifications. For the classification of hands vs. animals, we used a 2 (tDCS area: pMTG or mPFC) * 2 (training task: hands or tools) * 6 (hand-ROIs) ANOVA, whereas for the classification of tools vs. animals, we used a 2 (tDCS area: pMTG or mPFC) * 2 (training task: hands or tools) * 8 (tool-ROIs) ANOVA. We employed two separate ANOVAs because there were different ROIs per ANOVA.

As predicted, there was a significant interaction between tDCS area and training task such that the accuracies differed between the two tDCS areas for the classification between hands vs. animals (*F*(1,19) = 9.71, *p* = .006) and for the classification of tools vs. animals (*F*(1,19) = 6.88, *p* = .017). Specifically, post-hoc tests (FDR corrected; Benjamini & Hochberg, 1995) revealed that the accuracy for the classification of tool vs. animals was higher when tDCS was applied to pMTG and paired with the tool training task (pMTG_tools_), than when tDCS was applied to mPFC and paired with the tool training task (mPFC_tools_) (t(19) = 2.55, adjusted *p* = .04, see Figure 2C). However, there was no difference when tDCS was applied to pMTG in tandem with the hand training task (pMTG_hands_) nor when applied to mPFC in tandem with the hand training task (mPFC_hands_) (t(19) = 1.65, adjusted *p* = .12).

**Figure 2.**
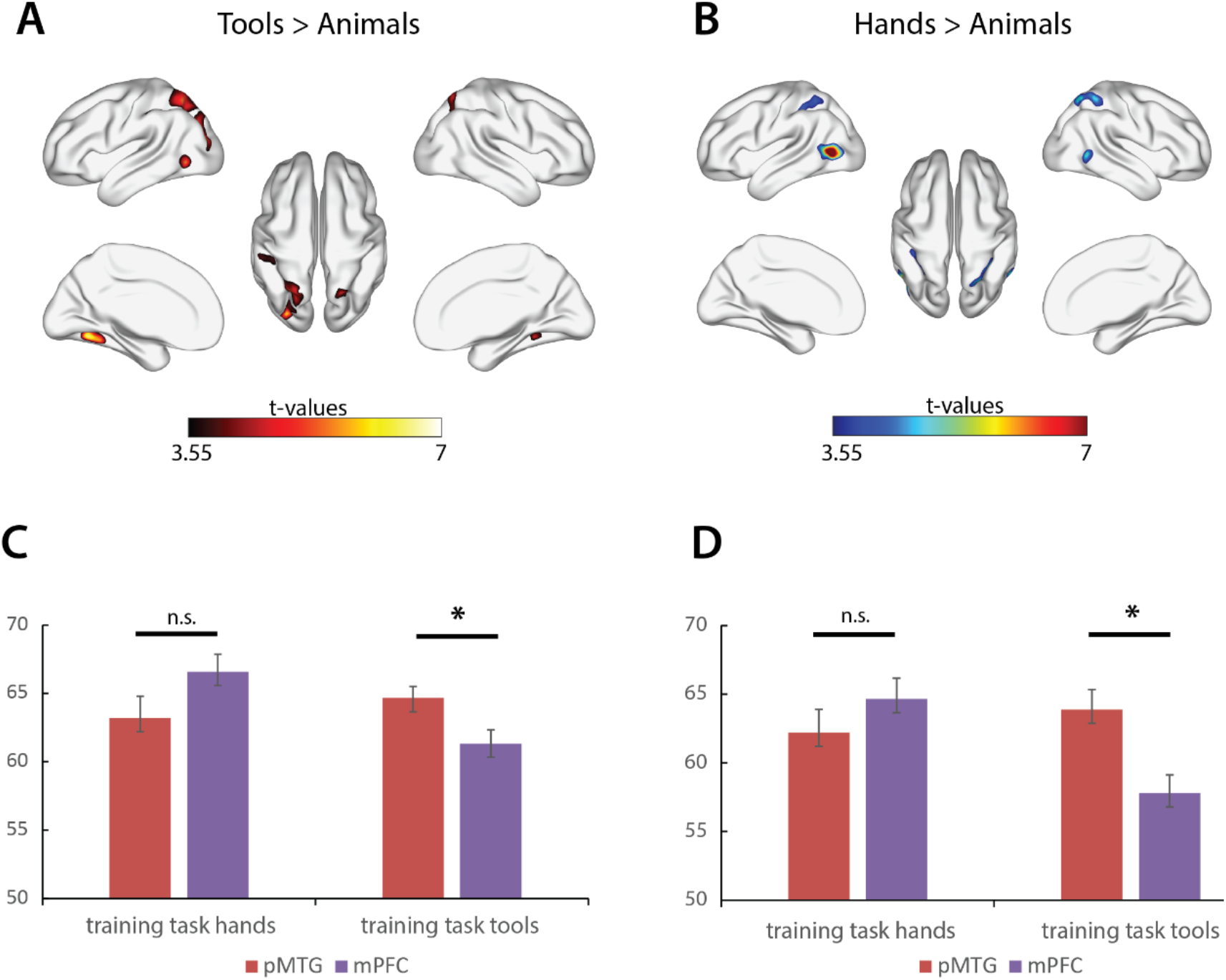
Contrasts of interest and MVPA results. The univariate results (p < .001, uncorrected) used to define regions of interest for the **(A)** tools > animals and **(B)** hands > animals contrasts; and classification accuracy (percentage) for **(C)** tools vs. animals and **(D)** hands vs. animals, for both tDCS areas (pMTG and mPFC) and training tasks (hands and tools), tDCS area demonstrating an interaction. All error bars reflect one standard error of the mean across participants (* = adjusted *p* value < .05).

Surprisingly, the accuracy for the classification hands vs. animals did not show the expected pattern. That is, hands vs. animals accuracy was higher for pMTG_tools_ than mPFC_tools_ (t(19) = 4.02, adjusted *p* = .002, see Figure 2D), while there was no difference between pMTG_hands_ and mPFC_hands_ (t(19) = 1.11, adjusted *p* = .28).

Because we already showed that the hand training task elicits significantly more errors and slower reaction times than the tool training task (see training task results), and that this training task did not show any significant difference for the two stimulation areas, we decided not to include this condition in the next analysis (i.e., we focused only on the tool training task; but see results for the hand task in Supplementary Figure 1).

In order to analyze the differences between stimulation sites for each ROI separately and see how stimulating pMTG concurrently with a tool training task changes classification accuracy, we compared the classifying accuracy for both classifications (tools vs. animals and hands vs. animals) between the conditions where we stimulated pMTG under a tool task (pMTG_tools_), and where we stimulated mPFC under a tool task (mPFC_tools_).

For the classification of tools vs. animals (Figure 3A and 3B), pMTG_tools_, when compared to mPFC_tools_, led to higher accuracy values only in two tool ROIs (and no hand ROIs): the left pMTG (t(19) = 4.28, adjusted *p* = .0008) and left mFUG (t(19) = 2.75, adjusted *p* = .05).

**Figure 3.**
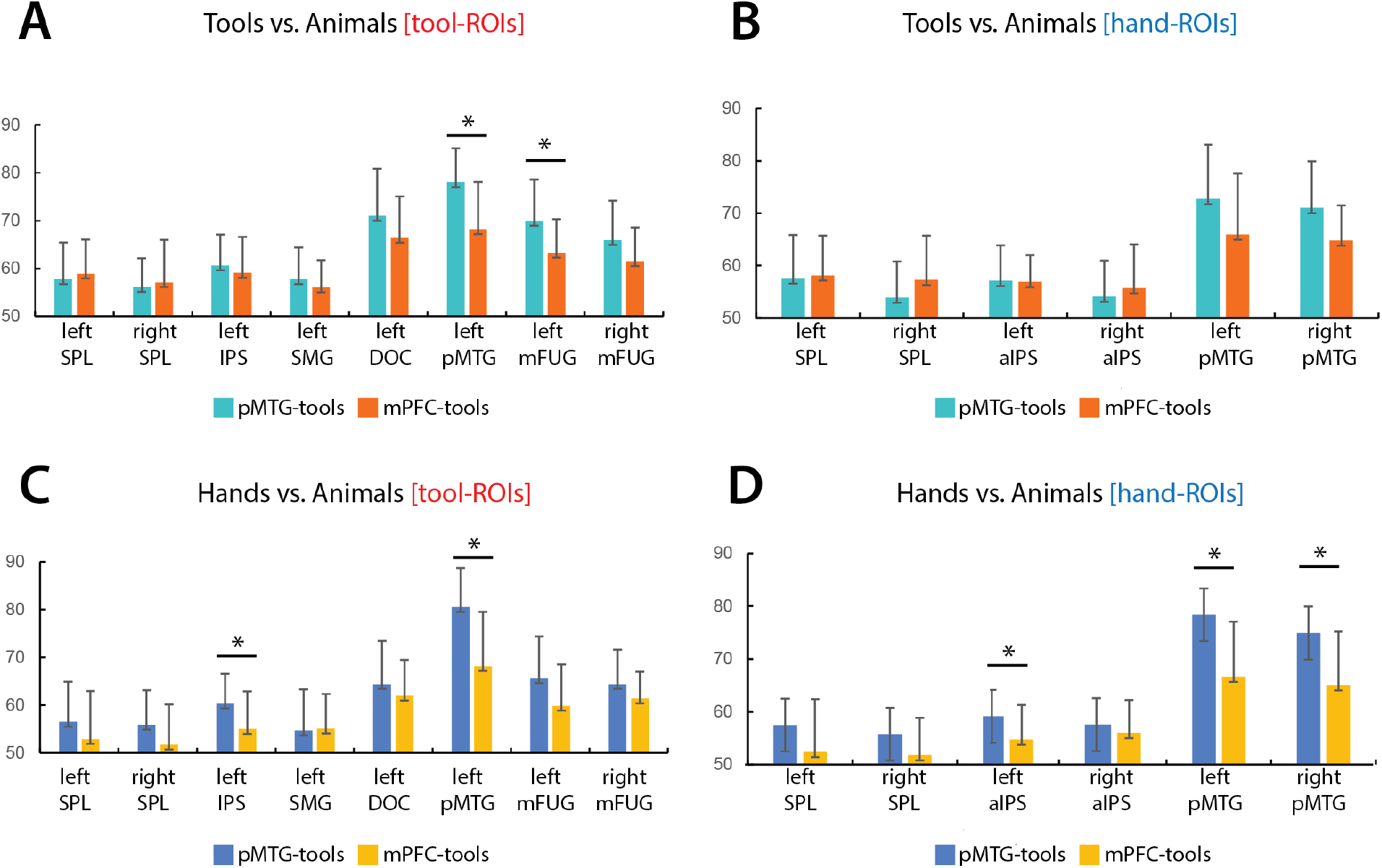
ROI-specific MVPA results. A comparison of the classification accuracy (percentage) between pMTG_tools_ and mPFC_tools_ for **(A)** tools vs. animals in each region identified as a tool ROI, **(B)** tools vs. animals in each region identified as a hand ROI, **(C)** hands vs. animals in each region identified as a tool ROI, and **(D)** hands vs. animals in each region identified as a hand ROI. P-values are FDR corrected for 8 tests in tool-ROIs and for 6 tests when analyzing hand-ROIs (* = adjusted *p* value < .05).

For the classification of hands vs. animals (Figure 3C and 3D), pMTG_tools_ (when compared to mPFC_tools_) led to significantly higher classification accuracies in three hand ROIs: left pMTG (t(19) = 5.72, adjusted *p* = .0006), right pMTG (t(19) = 3.66, adjusted *p* = .006) and left aIPS (t(19) = 2.78, adjusted *p* = .024). Moreover, it led to higher accuracy values in two tool ROIs and: tool -related regions: left pMTG (t(19) = 5.18, adjusted *p* = .0008) and left IPS (t(19) = 2.90, adjusted *p* = .036).

## Discussion

Here we investigated whether two functionally different networks that share certain nodes - as is the case with the tool and hand networks that share pMTG among other areas – could be disentangled by exploring horizontal modulations and long-distance connections through the use of tDCS. To do so, we stimulated pMTG, while combining it with category-specific training tasks in order to enhance the effects of the tDCS stimulation, and looked at how this affected classification accuracy between hands or tools (vs animals), when compared to a control stimulation site (mPFC). We showed that applying tDCS to an area where preferences for tools and hands overlap, such as pMTG (i) produced effects in distal brain regions, and (ii) partially facilitated the processing of categorical information in a way that was dependent on the training task prior and during tDCS stimulation.

Importantly, we were able to replicate previous studies (Lee et al., 2019; Ruttorf et al., 2019) by showing that tDCS stimulation modulates BOLD signal patterns in distal brain areas. Specifically, we demonstrated that classification accuracy for hands vs. animals and tools vs. animals was higher when pMTG was stimulated compared to mPFC in distal regions related to hand and tool processing. This result is in line with what previous studies showed (e.g., Lee et al., 2019; Ruttorf et al., 2019): object representations within a specific region can be causally modulated through horizontal modulations from a distal to a local region.

However, our results failed to fully confirm some of our predictions, especially for the training tasks that were coupled with tDCS. Here we predicted that there would be a category-specific effect of the task on the effects visible for the tool and hand networks – this prediction was not fully met.

On the one hand, we demonstrated that the classification between tools and animals benefited from the tDCS stimulation to pMTG only when this stimulation was paired with a tool training task, but not a hand training task, and only in tool ROIs. We showed that when classifying tools vs. animals, the pMTG_tools_ condition (when compared to the mPFC_tools_ condition), significantly improved the classification accuracy in left pMTG (as expected given that this was the tDCS stimulated area) and left mFUG. The effect on mFUG is an important one as it shows the importance of distal modulation on a functional network – stimulating a tool/hand overlap area under a tool training task led to an advantage in classifying tools vs. animals in a distal yet connected tool (but not hand) area – the left mFUG.

This is in line with our previous study (Amaral et al., 2021), where we showed that pMTG, when processing tools and in the process of conceptual integration, shares information with posterior parietal and dorsal occipital regions (associated to grasping - Almeida et al., 2008, 2010, 2014; Culham et al., 2003), and also communicates with mFUG (more related aspects of visual form and texture - Cant & Goodale, 2007; Cavina-Pratesi et al., 2010). Moreover, previous neuromodulation studies (Lee et al., 2019; Ruttorf et al., 2019) showed that interfering with the processing in a particular tool-region causally affects other distal regions of the tool-network. This suggests that despite the fact that pMTG is a tool/hand overlap region, it is possible to specifically target tool representations over hand representations, by combining the tDCS with a tool-task.

On the other hand, we were not able to obtain similar results for the hand training task over the classification of hands. Although this was an unexpected result, there may be some potential explanations for the failure to obtain results with the hand training task over hand classification. One possible explanation relates with the fact that the kind of task used for the hand training, unlike that for the tool training, was not necessarily related with recognition and processing of hands. Specifically, during the hand training task participants had to press a button every time the image changed from a right hand to a left hand (or vice-versa), whereas in the tool task, participants were instructed to look for a change in tool (e.g., from a hammer to a screwdriver). Thus, the task in the hand training condition could be more dependent on aspects related to mental rotation, rather than hand recognition and processing. In fact, as shown in the training result section, the hand task was clearly different from the tool task - accuracy during the tool task was above 90%, whereas it was around 75% for the hand task; reaction times for the hand task were, on average, about 300ms slower than for the tool training task. In part then, the lack of an effect for the hand training task may be related with the actual difficulty of the task, as well as its potential engagement of non-hand processes and networks. Moreover, the tDCS montage employed in this study, with the anodal electrode centered at pMTG and the two cathodal electrodes positioned both anterior and posterior to pMTG, may have inhibited the effect for the hand training task. The organization of the lateral occipitotemporal cortex (LOTC; Wurm et al., 2017) suggests that our tDCS montage, and specifically our most posterior cathodal electrode could have inhibited social and action representations important for the processing of hands (Bracci et al., 2010, 2018) and thus weakening, or completely overriding, the conjoint effect of the hand training and pMTG stimulation. These are two strong possible reasons for the lack of an effect of the hand tasks on the classification of hand stimuli, and we believe that future work will show that if these two aspects are taken care of, we will obtain a similar result for the hand training task as we did for the tool training task.

But our effects also show another potentially interesting but unexpected result – namely that the tool training task affected hand classification in certain tool and hand ROIs. In particular the areas where we show an effect of the tool training task on hand classification are areas typically associated with object grasping and manipulation, and object-related action (i.e., pMTG and left aIPS). For instance, aIPS is known to play an important role in the computation of hand-shapes for object grasping (Binkofski et al., 1998, 1999; Culham et al., 2003; Monaco et al., 2011), particularly in shaping the hand for the correct manipulation of the object (Buchwald et al., 2018). Perhaps then, the tool training task leads to an advantage for hand classification because it is distally engaging regions dedicated to the processing of grasping and manipulation properties. For instance, grasping a tool requires information about object structure and object volumetry (e.g., Brandi et al., 2014; Buxbaum et al., 2007) – when we want to manipulate a hammer, we need to adjust our hand based on the handle format and size. Thus, it is possible that the tool training task led to tool-related distal activations in grasping, manipulation and action areas, and by virtue of that, and of the fact that hand processing in those areas was intimately related with action, those distal tool-related effects percolated to hand processing, and hence hand classification. Moreover, the hand images used during our fMRI experiment were actually in grasping postures (power and precision grips) and this could produce an activation of the motor system. In fact, we have previously demonstrated shared tool-hand invariant (power vs. precision) grasp-type representations in the left posterior parietal cortex (Bergström et al., 2021). Additionally, several studies have shown that pictures of hand grasp postures can influence object categorization, such that visualizing a hand with a particular grasp posture activates motor information, affecting the processing of manipulable objects, like tools (e.g., Almeida et al., 2018; Borghi et al., 2007; Craighero et al., 2002).

Overall, then, our results (at least partially) show that overlapping functionally-specific networks can be disentangled by focusing on their category-specific horizontal modulations between neural nodes. Specifically, if we focus on how these horizontal long-distance modulations causally affect local processing, we will bring forth strong category-specific organizational dissociations.

## Acknowledgements/Funding

This work was supported by the Portuguese Foundation for Science and Technology (R&D grant PTDC/PSI-GER/30745/2017 and fellowship CEECIND/03661/ 2017 to F.B., doctorate scholarship SFRH/BD/114811/2016 to L.A., doctorate scholarship SFRH/BD/137737/2018 to D.V.); the European Research Council (ERC) under the European Union’s Horizon 2020 research and innovation programme (Starting Grant number802553 “ContentMAP” to J.A); the University of Padova, Department of Psychology and Human Inspired Technology Centre in Padova (doctorate scholarship to R.D.). R.D. carried out the present work within the scope of the project “Use-inspired basic research”, for which the Department of General Psychology of the University of Padova has been recognized as “Dipartimento di Eccellenza” by the Ministry of University and Research).

---

**Supplementary Figure 1.**
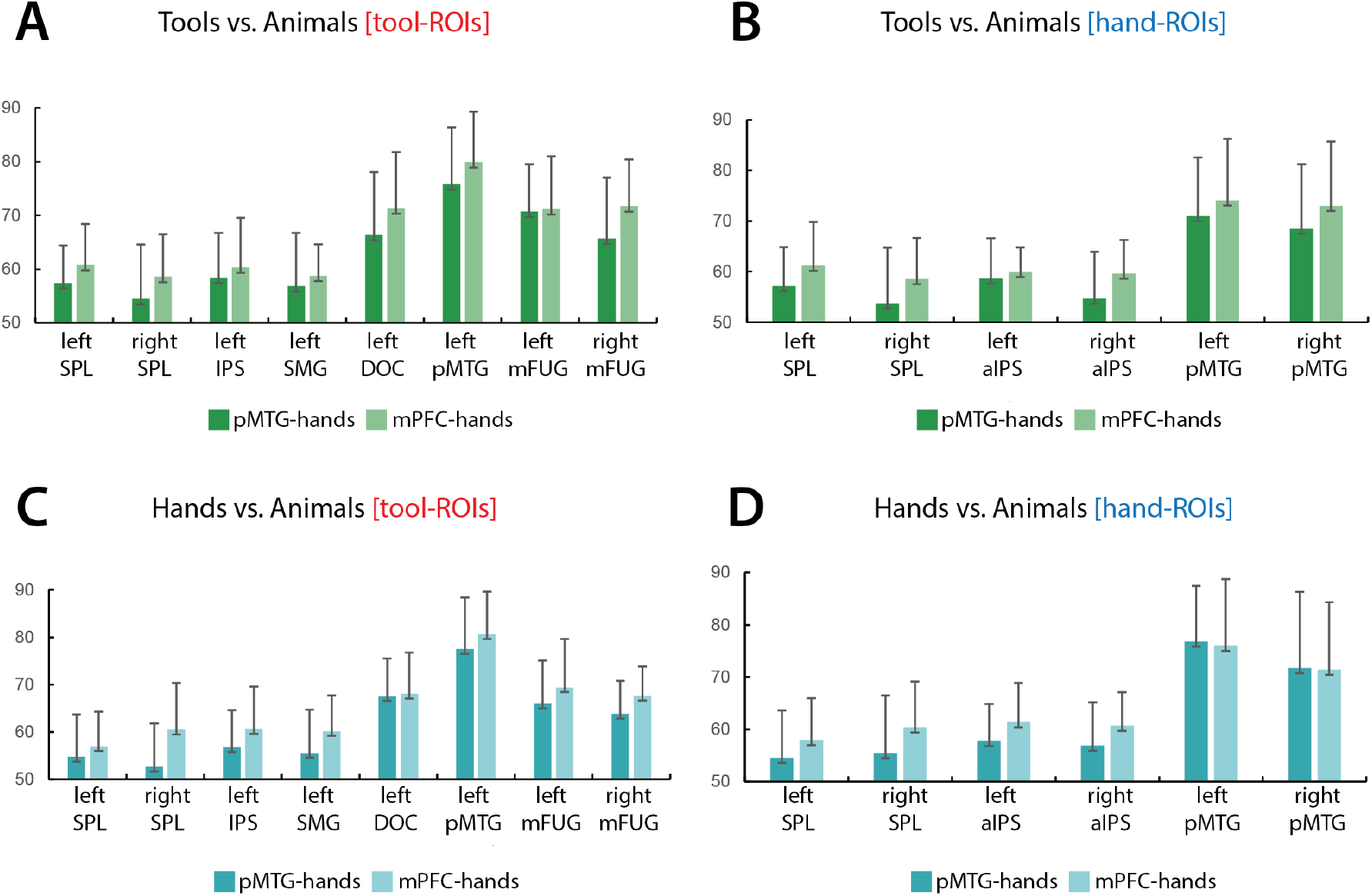
ROI-specific MVPA results (hands training session). A comparison of the classification accuracy (percentage) between pMTG_hands_ and mPFC_hands_ for **(A)** tools vs. animals in each region identified as a tool ROI, **(B)** tools vs. animals in each region identified as a hand ROI, **(C)** hands vs. animals in each region identified as a tool ROI, and **(D)** hands vs. animals in each region identified as a hand ROI. P-values are FDR corrected for 8 tests in tool-ROIs and for 6 tests when analyzing hand-ROIs and show no significant results (all adjusted *p* values > .1).

